# Compound-SNE: Comparative alignment of t-SNEs for multiple single-cell omics data visualisation

**DOI:** 10.1101/2024.02.29.582536

**Authors:** Colin G. Cess, Laleh Haghverdi

## Abstract

One of the first steps in single-cell omics data analysis is visualization, which allows researchers to see how well-separated cell-types are from each other. When visualizing multiple datasets at once, data integration/batch correction methods are used to merge the datasets. While needed for downstream analyses, these methods modify features space (e.g. gene expression)/PCA space in order to mix cell-types between batches as well as possible. This obscures sample-specific features and breaks down local embedding structures that can be seen when a sample is embedded alone. Therefore, in order to improve in visual comparisons between large numbers of samples, we introduce Compound-SNE, which performs what we term a soft alignment of samples in embedding space. We show that Compound-SNE is able to align cell-types in embedding space across samples and data modalities, while preserving local embedding structures from when samples are embedded independently.

## 1 Introduction

Visualization of high-dimensional data is a key aspect when examining single-cell omics (epigenomis, transcriptomics, proteomics, etc.) data samples. Many different algorithms exist for embedding high-dimensional data into two-dimensional space, though t-SNE (t-distributed Stochastic Neighbours Embedding) and UMAP (uniform manifold approximation and projection) remain the most common (1; 2). Besides visualizing a single sample, it is important to be able to visually compare multiple single-cell samples, e.g. scRNA-seq from different patient samples or data modalities such as paired scRNA-seq and scATAC-seq data (3) on the same sample or same patient, (multi-view data) before moving on to further analyses. If the dataset is complete (i.e., containing all cell states of interest) the reference dataset can be first embedded and other data sets projected onto it (4; 5; 6). Otherwise, in current approaches, data integration is performed to merge samples together, which are then embedded all at once (7; 8; 9). While this does achieve a good alignment of different samples, data integration algorithms modify gene expression values in order to best mix samples together, leading, in embedding space, to the dissolution of unique local structures that are seen in original, unintegrated embeddings. Although data integration is still important for other analyses (e.g. such as cell type label transfer tasks (10)), we propose here an alternative method for visualizing multiple single-cell samples. Compound-SNE performs what we term a soft alignment, aiming to maximize the alignment of multiple embeddings while minimizing the local structural differences from the samples’ independent embeddings. This is done in a two-step process: (1) alignment in PCA space via matrix transformation in order to align embedding initializations, and (2) addition of a force term to the embedding algorithm, which pulls clusters of cells together based on annotations.

## 2 Alignment Overview

The complete workflow of Compound-SNE consists of five steps as follows. Compound-SNE is designed to work with Scanpy (11) formatting, taking in an AnnData object. Compound-SNE is available at https://github.com/HaghverdiLab/Compound-SNE

### 1. Data processing

Data should be processed via whatever method the user deems suitable and transformed into PCA (principal component analysis) space. Following Scanpy, this should be stored in the AnnData object as .obsm[‘X_pca’]. While different samples and modalities may contain different number of original features, they should share the same number of features in PCA space, though data integration should not be performed here.

Additionally, for alignment, Compound-SNE requires cell annotations, ideally cell types, though other types of annotations are acceptable. If this is not available, Compound-SNE takes one sample as a reference, performs k-means clustering, and, using the cells closest to each cluster centroid, identifies corresponding centroids in the other samples using mutual nearest neighbors (7) in PCA space. We note that this in not preferred and that these clusters are only used for visual alignment and not any type of functional identification. Compound-SNE then integer encodes annotations. For the rest of the paper, we will refer to annotations as cell types.

### 2. Reference selection

Compound-SNE requires at least one sample to use as a reference. If not specified, Compound-SNE first chooses a primary reference as the sample with the most unique cell types. This sample is used for the primary alignment, as described in the following section. Then, if the primary does not contain all of the cell types, secondary references are chosen in order to complete the set of cells.

### 3. Primary alignment

Samples are first aligned in PCA space via matrix transformations in order to align cell-type centers. A cell-type center (only for shared cell types) x components matrix is found for each sample, which is then aligned to the primary reference, after scaling, via a Procrustes transformation (12), which scales and rotates a matrix to minimize the sum of squared errors from a reference matrix. The obtained transformation matrix is used to transform the full sample.

### 4. Embedding initialization

The embedding portion of Compound-SNE is based on the openTSNE python package. As (13) shows, initialising a non-linear embedding optimization with PCA components enhances preservation of global structures. The authors also report that both t-SNE and UMAP equally preserve global structures when using the same initialisation. We therefore use the first two components of the transformed PCA space in order to initialize the embedding for each sample.

### 5. Alignment via forces

To obtain better alignment between samples with minimal disturbance to local embedding structure, we include an additional force term to the embedding process that pulls the centers of cell type clusters (that may deviate from the primary alignment in the process of t-SNE iterations) together for each sample. We first embed the reference sample as normal, then find the centers of each type in embedding space. When embedding the remaining samples, during each embedding step, type centers are found and unit vectors in the direction of the respective reference sample centers are calculated. These vectors are multiplied by a scalar force value and applied to each cell of the respective type. This means that cells of the same type move in parallel in the direction of the reference embedding, thus not disturbing cluster shape. For the practical implementation, we calculate this force in between each iteration of openTSNE.

Because not all samples may contain every cell type, as described above, the primary reference is chosen as the one with the most unique type. We then identify secondary references, using the minimum needed to create a set containing all of the present cell types. Secondary references are then aligned sequentially to the primary, using their embeddings to obtain embedding centers of remaining type.

## 3 Application

We apply Compound-SNE to datasets consisting of multiple patients and modalities, demonstrating its utility for comparing different but related datasets. One dataset consists of bone marrow samples from six healthy patients, containing both gene expression and surface markers (14). The second dataset consists of kidney samples from five healthy patients, both gene expression and ATAC (15). A subset of alignments is shown in Figure 1, with full alignments in Figure S1.

**Figure 1:**
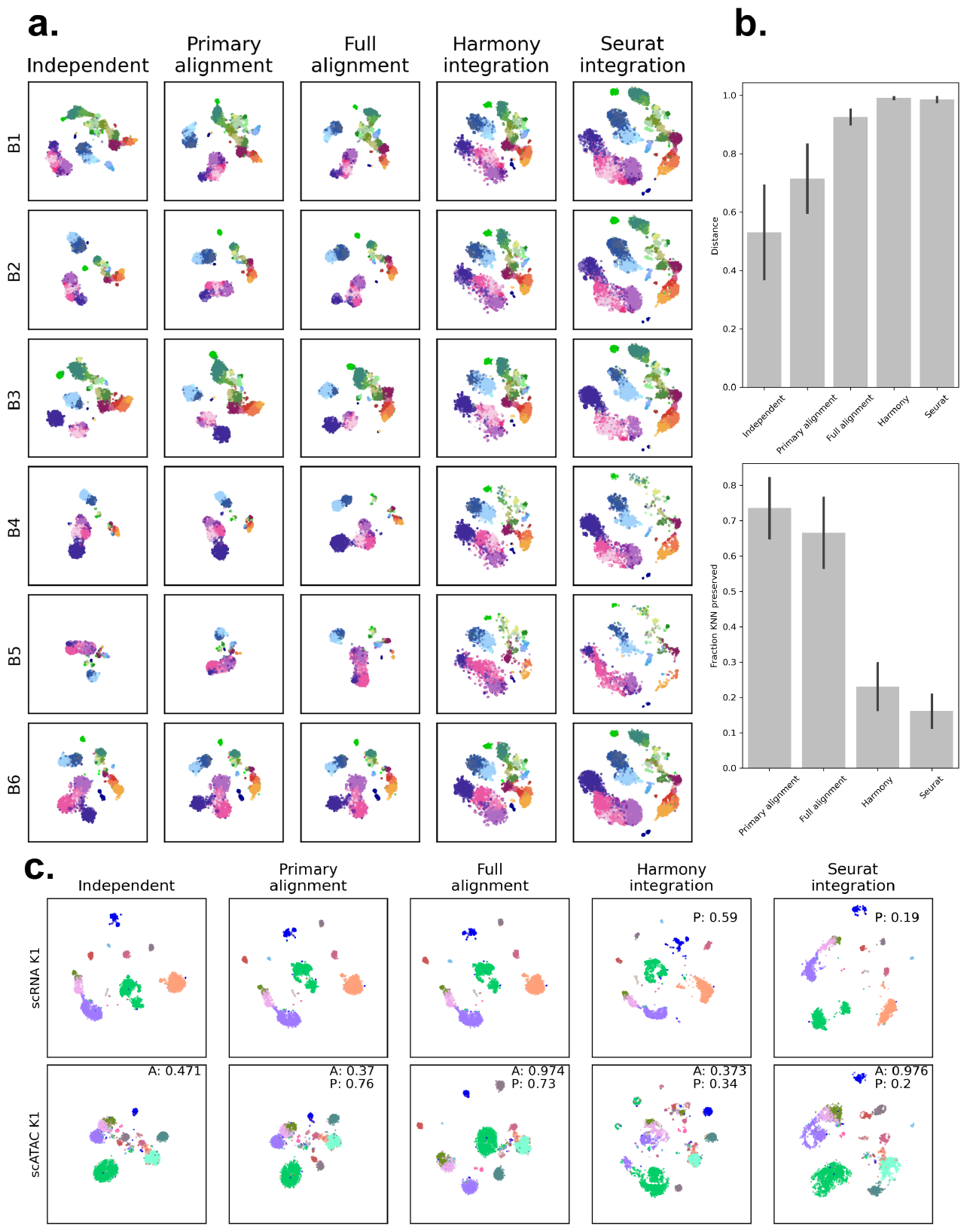
(a) Embeddings for the six patients from the bone marrow dataset, using B6 as the primary reference. Each row corresponds to a different alignment/integration method. All embeddings are on the same spatial scale. (b) Metrics for the embeddings shown in A. Means with error bars for standard deviation. Top: structure preservation, calculated as the fraction of KNN for each point preserved from the Independent embeddings. Bottom: alignment of the embeddings as the distance between normalized cell type centers. (c) Alignment/integration of scRNA and scATAC samples for K1 of the kidney dataset. Alignment scores of scATAC to scRNA are shown on each scATAC subplot, labeled as A. Structure preservation scores, labeld as P, for scATAC are shown on the subplots, excluding the Independent embedding. This score is also shown for integration methods on scRNA. All embeddings coordinates are on the same scale. Cell-type legends for (a) and (b) are shown in Figure S5

In Figure 1a, using gene expression of patient B6 as a reference, we show that Compound-SNE can be used to align gene expression for several patients. The first column shows the original, independent embeddings for each sample. The second columns shows embedding following the primary alignment and the third column shows embedding with the additional force term. The final two columns shows embedding following data integration using Harmony and Seurat, as a comparison to our method. Visually, we see that even using only the primary alignment offers a reasonable improvement over the independent embeddings, with the full alignment providing a much greater visual alignment. Notably, the full alignment yields embeddings that retain much of the cluster shapes that are seen in the independent embeddings. The two integration methods, while clearly aligning all of the samples, visually erase much of the structures unique to each patient in the independent embeddings. This is because cells are forced to mix well between batches.

In Figure 1c, we align scRNA and scATAC samples from the same patient. While in comparison to Figure 1a, where the independent embeddings look somewhat comparable between patients (as well as between scRNA and surface markers in Figure S1), the embeddings for scRNA and scATAC look very different from each other initially, obscuring comparison. Primary alignment achieves a modest improvement, while the full alignment yields a much stronger improvement while preserving original cluster shapes. We were unable to integrate the two modalities using Harmony, while Seurat was able to integrate them, again at the cost of dissolving structures present in the independent embeddings.

## 4 Comparison statistics and evaluations

Beyond a visual comparison of embeddings, we calculate several metrics to compare how well-aligned embeddings are to each other and how well embedding structures are preserved between aligned embeddings and the original embeddings.

### 1. Alignment score

Beyond visually comparing embeddings, we calculate a metric to determine how well-aligned samples are. In embedding space, we find the centers of each celltype for each sample and take the sum of squared errors between points. This value, d, is then transformed via 1/(1 + d) so that a value closer to 1 indicates a better alignment. We see that (Figure 1b, top), as we progress from independent embeddings to aligned initializations to aligned with center-based force, we get better alignment, which is consistent with the visual results. We do see that data integration methods Harmony and Seurat yield the best alignment between samples, which is expected based on the nature of data integration. Alignment scores between scRNA and scATAC for patient K1 are shown directly on the plots of Figure 1c.

### 2. Locality preservation

While data integration yields the best alignment between samples, we can visually see that this is at the cost of the original embedding structure (Figure 1b, bottom). To determine the preservation of local structures present in each embedding, we calculate the k nearest neighbors for each cell in the independent embedding and compare it to the nearest neighbors in each alignment, taking the fraction shared as a metric of structure preservation. We see that the primary alignment obtains the best preservation of original structure, with alignment with center-forces performing only slightly worse. Data integration, on the other hand, greatly disrupts these local structures. We therefore see that there is a trade-off between structure preservation and sample alignment. Preservation scores for scRNA and scATAC for patient K1 are shown directly on the plots of Figure 1c.

### 3. Alignment of data views with highly variable sizes (cell numbers)

Furthermore, to demonstrate the alignment of samples with highly different cell densities, we randomly subsample bone marrow B2 to 696 cells (1/10 of the cells) and align it with the full sample for B1 (9751 cells) (Figure S2). We see that this still achieves a nice visual alignment.

### 4. Computational efficiency

In Figure S3, we compare the runtime for each samples when embedded independently and embedded with alignment forces. We find that, with a couple of outliers in either direction, the addition of alignment forces does not impact runtime. As expected, smaller datasets tend to embed faster.

### 5. Clustering for cell annotations

We mentioned that Compound-SNE is able to generate non-cell-type-specific annotations for the sake of performing alignment.

Applying the full alignment to these generated annotations for the bone-marrow scRNA samples is shown in Figure S4, which shows a comparable alignment, in this case, to using the original cell-type annotations.

## 5 Conclusion

With Compound-SNE, we demonstrate how we can perform a soft alignment of embeddings for single-cell samples from different patients and modalities. This aids a visual comparison between many samples, with minimal disturbance to the unique sample structures seen when embedding samples independently.

