## Supplementary Figures for "Compound-SNE: Comparative alignment of t-SNEs for multiple single-cell omics data visualisation"

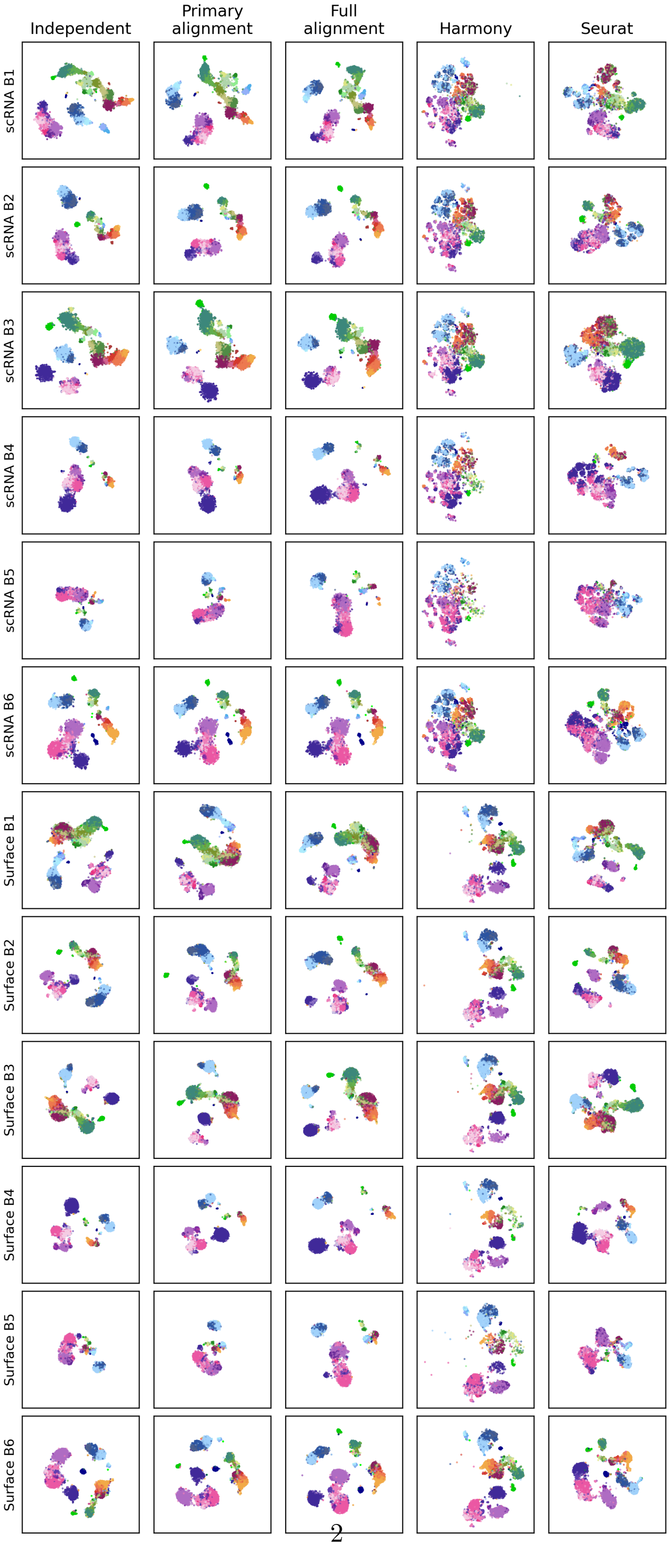




Figure S1: Alignment of scRNA and surface markers for all bone marrow patients

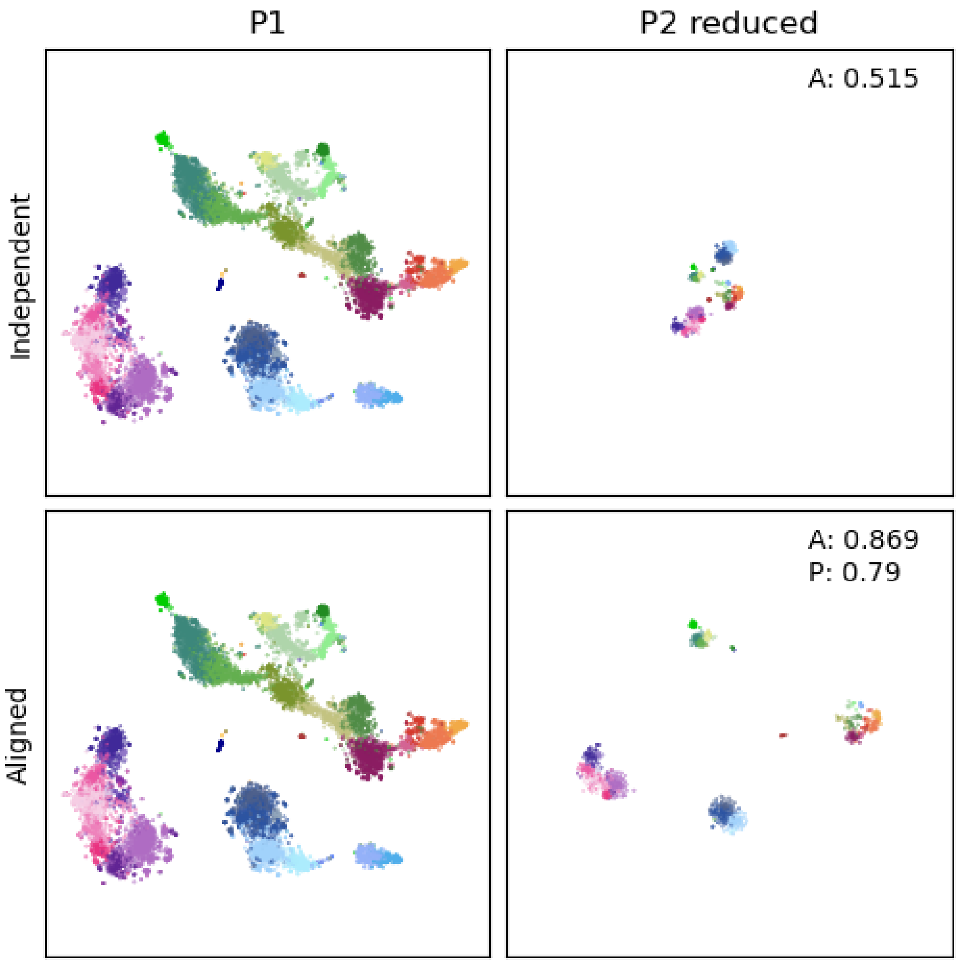


Figure S2: Alignment of scRNA for B1 and B2, with B2 randomly subsampled to 1/10 the size. All embeddings are shown on the same spatial scale.

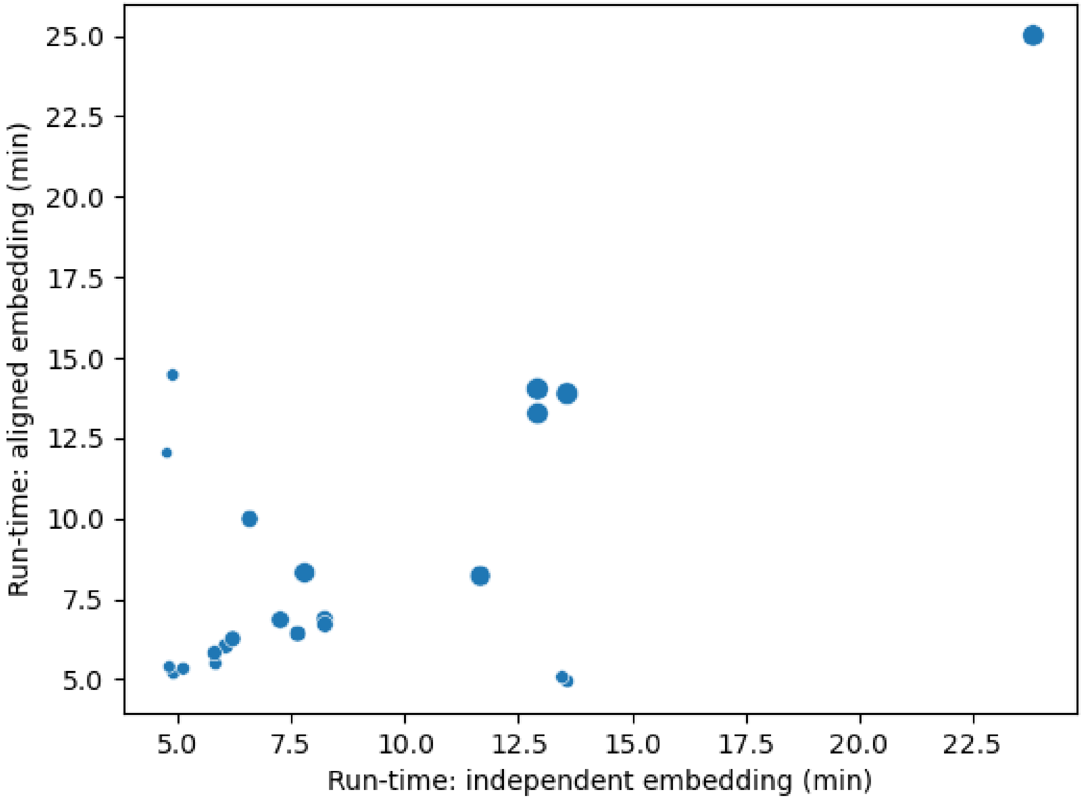


Figure S3: Runtime for independent and aligned embeddings. Each dot represents one sample. Dot size corresponds to sample size.

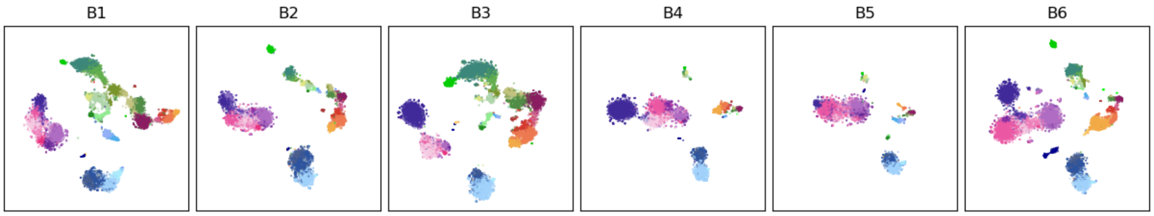


Figure S4: Alignment via kmeans clustering and MNN.

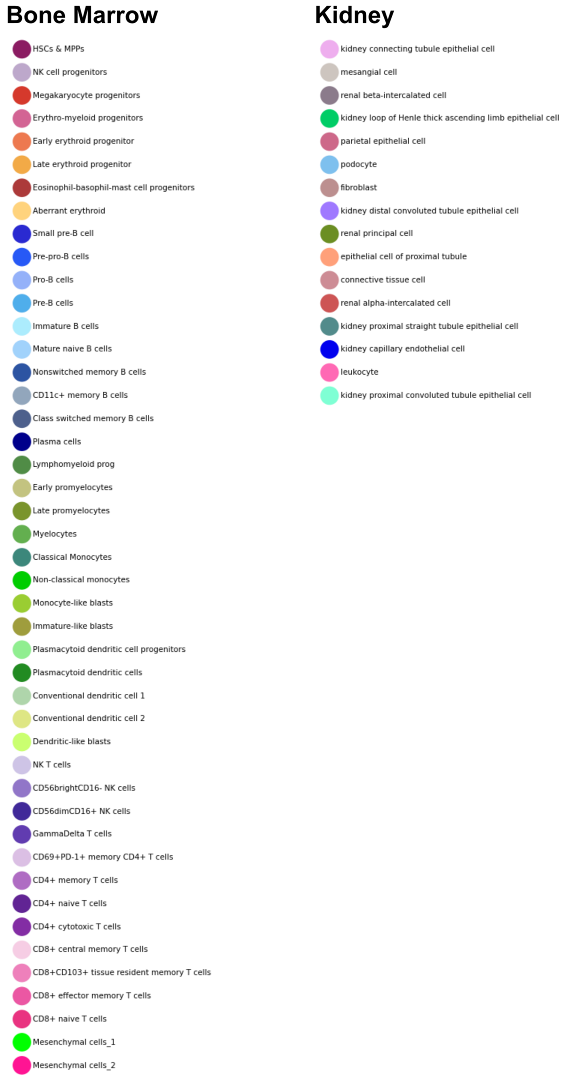


Figure S5: Cell-type legends for the two example datasets.
